# Genomic Insights into a Highly Specific Marine Symbiosis Uncovers Geographic Structuring Without Co-Divergence Between *Siphamia* Cardinalfish and Their Bioluminescent Symbiont

**DOI:** 10.1101/2025.05.30.657134

**Authors:** Emily Neff, Alison Gould

**Affiliations:** Institute for Biodiversity Science and Sustainability, California Academy of Sciences, San Francisco, CA 94121, USA; College of Science and Technology, Department of Biology, Temple University, Philadelphia, PA 19122, USA

## Abstract

Symbiotic relationships with microorganisms are fundamental to life on Earth, yet relatively little is known about how these interactions persist through time, how they co-evolve, and to which degree they are genetically constrained. In this study, three cardinalfish species in the genus *Siphamia, S. tubifer, S. mossambica*, and *S. fuscolineata*, were analyzed for patterns of genetic divergence along with their luminous bacterial symbionts, from locations throughout the hosts’ broad Indo-Pacific distribution. Using whole genome sequencing (WGS) of the fish light organ, we investigated whether the specificity of the association is maintained across host species and over this geographic range and whether there are any patterns of symbiont divergence associated with either. Our results indicated that the light organ symbionts of all three *Siphamia* species examined were identified as *Photobacterium mandapamensis*, suggesting a high degree of specificity of the symbiosis is conserved across hosts and geography. There was evidence of biogeographic structure in the symbiont between the three sampling regions, but no co-diversification between the hosts and their symbionts (p = 0.92). However, an analysis of single nucleotide polymorphisms (SNPs) between two *S. tubifer* populations sampled from Japan and the Philippines indicated some genetic differentiation (F_ST_ = 0.043) with phylogenetically-distinct clades of symbionts. Overall, these findings indicate that the association between *Siphamia* hosts and *P. mandapamensis* is highly conserved, yet there is significant genetic diversity within the symbionts driven by geography and potentially host ecology.

## BACKGROUND AND SIGNIFICANCE

Microbial symbioses are an essential part of nature, forming intricate partnerships that shape ecological communities, influence host fitness, and drive evolutionary processes. Among these associations, bioluminescent symbioses represent fascinating examples where host animals gain the ability to produce light through a symbiotic relationship with luminous bacteria. These light-producing symbioses have independently evolved multiple times across diverse marine lineages, enabling novel ecological functions such as predator avoidance, prey attraction, and intraspecific communication (Lau & Oakley 2021; Davis et al. 2016). Despite their ecological importance and widespread occurrence, our understanding of the genetic mechanisms governing host-symbiont specificity, symbiont transmission, and co-evolutionary dynamics within bioluminescent systems remains limited, particularly across geographic scales and among closely related host species. The bioluminescent association between cardinalfish in the genus *Siphamia* and their bacterial partner *Photobacterium mandapamensis* offers an ideal opportunity to address these fundamental questions in microbial symbiosis.

The association between cardinalfish in the *Siphamia* genus and their bioluminescent symbiont, *Photobacterium mandapamensis* is an emerging model for studying strain-level genetic diversity in the symbiosis (Gould & Osland 2025). There are currently 25 described species of *Siphamia* located throughout the Indo-Pacific (Gon & Allen 2012). A unique characteristic of the genus is a disk-shaped light organ attached to the intestine that houses the bioluminescent symbiont (Dunlap & Nakamura 2011). The bacterially-produce light is believed to be used for countershading while foraging at night. Within *Siphamia*, there are two distinct light organ morphologies, which aid in classifying *Siphamia* into groups: *S. tubifer* and *S. tubulata* (Gon & Allen 2012), herein referred to as the ‘*tubifer’* and ‘*tubulata’* groups, respectively. The light organ and its associated tissue in the *tubifer* group has a striated pattern, while that of the *tubulata* group has a dotted pattern (Gon & Allen 2012).

*Siphamia tubifer*, a member and namesake of the *tubifer* group, has the largest geographic distribution of all *Siphamia* species (Gon & Allen 2012). Consequently, it has also been the most widely studied species (Leis and Bullock 1986; Dunlap et al. 2011; Gould et al. 2015, Gould and Osland 2025). Most members of the *tubifer* group are morphologically similar displaying two color patterns — silver and black striped or dark brownish black, with an average standard length between 2-3 cm (Gon & Allen 2012). These fish typically associate with venomous echinoderms, such as sea urchins and the crown-of-thorns sea star, which provide protective shelters throughout the day. Many species within the *tubifer* group exhibit overlapping ranges (Gon & Allen 2012).

*Siphamia* hosts acquire their symbiont while they are in their pelagic phase during larval development. Male fish brood the eggs in their mouth for 4-5 days, after which the larvae are released into the water column, where they continue to develop until settlement on their chosen substrate approximately 30 days post-hatch (dph) (Gould et al. 2016). Once the bioluminescent symbiont is ingested, it enters the light organ through a duct connected to the fish’s intestine. The light organ is comprised of cube-shaped epithelial cells that form chambers where the bacteria is housed. To regulate the symbiont population, the host regularly sheds symbiont cells through the duct and back into the environment with fecal waste (Dunlap & Nakamura 2011).

While many species of bioluminescent bacteria form facultative partnerships with marine fish and squid (Kaeding et al. 2007; Nyholm and McFall-Ngai 2021), *Siphamia* seem to exclusively associate with *P. mandapamensis* (Kaeding et al. 2007; Gould et al. 2021), a subspecies of *P. leiognathi* (Urbanczyk et al. 2010). However, the symbiosis has primarily been characterized for one *Siphamia* species, *S. tubifer*, within a relatively small geographic region in the Okinawan Islands, Japan (Gould & Dunlap 2019) as well as for a handful of museum specimens with limited genetic data (Gould et al. 2021). Despite overlapping geographic ranges and shared morphological and ecological traits, the diversity, specificity and evolutionary patterns of the bioluminescent symbionts of the other *Siphamia* hosts in the *tubifer* group remain unexamined.

This study investigates whether the high degree of specificity with *P. mandapamensis* is maintained across host species within the *tubifer* group sampled over a broad geographic area and whether there were any notable divergence patterns in the association. We examined three *Siphamia* species—*S. tubifer, S. fuscolineata*, and *S. mossambica*—from four locations: Okinawa, Japan; Caubyan Island, Philippines; Verde Island, Philippines; and Zanzibar, Tanzania. We first characterized symbionts of the various hosts to determine whether the specificity with *P. mandapamensis* is maintained. Using genome-wide data, we then analyzed patterns of genetic variation in the host and symbiont and tested for any evidence of co-divergence. In doing so, we also characterized the divergence patterns of *Siphamia* species with very similar life histories while helping to resolve the evolutionary history of the *tubifer* group of *Siphamia*.

## METHODS

### Specimen Collection and Light Organ Extraction

A total of 61 *Siphamia* specimens were collected from four locations throughout the Indo-Pacific including Okinawa, Japan (n=15), Caubyan Island, Philippines (n=10), Verde Island, Philippines (n=24), and Zanzibar, Tanzania (n=12) (Supplementary Figure 1). Most specimens were collected at a depth of approximately 3 meters, with the exception of the Verde Island individuals, which were collected at approximately 30 meters depth. Specimens were collected offshore using hand nets and euthanized following an approved IACUC protocol (2021-12 IMSVBS) at the California Academy of Sciences. All specimens were kept at -80°C until further processing.

Each fish specimen’s light organ was aseptically dissected, and its total genomic DNA was extracted following the QIAGEN DNeasy Blood and Tissue Kit (QIAGEN) Quick-Start protocol. A Nanodrop 2000C Spectrophotometer (Thermo Fisher Scientific) was used to assess DNA purity, and a Qubit dsDNA HS Assay Kit (Thermo Fisher Scientific) was used to measure the DNA concentration on a Qubit 4.0 Fluorometer (Invitrogen). All extracted DNA was stored at -20°C until library preparation.

Library preparation was carried out on the 61 total DNA samples using the NEBNext Ultra II FS DNA Library Prep kit (New England Biolabs) for whole genome sequencing. Samples were individually indexed using the NEBNext Multiplex Oligos for Illumina (New England Biolabs). After library preparation, the DNA concentration was measured again using a Qubit 4.0 fluorometer (Invitrogen), and samples were normalized to 2ng/μL. The molarity was estimated using a 4150 TapeStation system (Agilent Technologies) for pooling and sequenced individually as paired-end 150 bp reads using the Novaseq 6000 Platform (Illumina) by Novogene Corporation Inc. (Sacramento, CA).

Raw sequence reads were filtered and trimmed using fastp v0.22.0 (Chen et al. 2018; Chen 2023) for paired-end data with the flag -l for both fish and bacterial DNA. Two reference genomes were used for aligning the fish and bacteria reads separately with BWA-MEM v0.7.17 (Li 2013); the host reference is *Siphamia tubifer* (ASM2046626v1, Gould et al. 2022) and the bacterial reference is *Photobacterium mandapamensis Ik8*.*2* (ASM3068539v1, Gould & Henderson, 2023).

### Host Analyses

After quality filtering, four individuals were removed from the analysis. Filtered sequences from the remaining 57 individuals were first aligned to the whole *S. tubifer* mitochondrial genome (Gould et al. 2022) using BWA-MEM v0.7.17. The aligned reads were then sorted and indexed, and consensus FASTA files were generated for each individual using SAMtools v1.19 (Danecek et al. 2021), which were used to create a multi-alignment using AliView v1.28 (Larsson 2014). Three additional cardinalfish species were included in the analysis as references: *Apogonicthyoides taeniatus* (MN699562.1), *Ostorhinchus fasciatus* (NC_058293.1), and *Sphaeramia orbicularis* (AP018927.1). A phylogeny was then inferred in IQ-TREE v2.0.3 (Nguyen et al. 2014) using the TPM2+F+R5 model based on the BIC score with SH-aLRT and ultra-fast bootstrap support based on 1000 replicates.

An additional analysis of the cytochrome oxidase I (COI) gene was carried out to infer the phylogenetic relationships among *Siphamia* species for which sequence data was available from NCBI. Sequences of the 57 hosts were aligned to the *S. tubifer* COI gene using BWA-MEM v0.7.17, SAMtools v1.19, and AliView v1.28. An additional 31 COI sequences from *Siphamia* species along with the reference cardinalfish species from the whole mitogenome analysis were included in the analysis. The phylogeny was inferred using IQ-TREE v2.0.3 with the substitution model TPM2+F+I+G4 based on the lowest BIC score with 1000 SH-aLRT and ultra-fast bootstrap replicates.

To analyze patterns of genetic differentiation in the host, reads that aligned to the *S. tubifer* reference genome were filtered and sorted using SAMtools v1.19. The paired reads were also deduplicated using MarkDuplicates Picard v3.1.0 (https://broadinstitute.github.io/picard/). Single nucleotide polymorphisms (SNPs) were identified using the GATK v4.5.0.0-0 (McKenna et al. 2010; Van der Auwera & O’Connor 2020) best practices workflow for whole genomes (https://gatk.broadinstitute.org/hc/en-us/sections/360007226651-Best-Practices-Workflows). SNPs were called and genotyped based on chromosome position, and were quality assessed based on allele frequency (MAF ≥ 0.05), mean depth per individual (minDP = 8, maxDP = 60), site quality (minQ > 30), and proportion of missing genotype data (max-missing ≥ 95%), using VCFtools v0.1.16 (Danecek et al. 2011) for each individual. All 23 chromosomes were concatenated into one GVCF file using BCFtools v1.19 (Danecek et al. 2021). Analyses were performed on SNPs at both the species level, between *S. tubifer, S. mossambica*, and *S. fuscolineata*, and at the population level, between the two *S. tubifer* populations.

Sequence data from 57 *Siphamia* individuals were used to perform a principal component analysis (PCA) across species based on 15,506 SNPs from the post-filtered GVCF file. The R package adegenet v2.1.10 (Jombart & Ahmed 2011) was used to compute the scaled allele frequencies, and ade4 v1.7-22 (Bougeard & Dray 2018) was implemented to generate PCA plots. A permutational multivariate analysis of variance (PERMANOVA) was applied based on distance matrices using vegan v.2.6-8 (Anderson 2001; Warton et al. 2012).

An additional PCA was carried out using the same approach on 1,098,423 SNPs identified from the 23 *S. tubifer* specimens from the Philippines and Japan from the post-filtered GVCF files to examine patterns of genetic variation between individuals from these two locations. Another PERMANOVA was applied at the population level. To identify outlier loci that are putatively under selection, an F_ST_ analysis between the two populations was performed using the hierfstat package v0.04-22 (Goudet 2004). Additionally, observed and expected heterozygosity, genotype concordance, and allele frequency were calculated for each population using adegenet v2.1.10 and dplyr v1.1.4 (Wickham et al. 2023). Nucleotide identity was calculated using R packages vcfR v1.15.0 (Knaus and Grünwald 2016; Knaus and Grünwald 2017), ape v5.7-1 (Paradis & Schliep 2019), and pegas v1.3 (Paradis 2010). The R package qqman v0.1.9 (Turner et al. 2014) was used to create a Manhattan plot to visualize putative SNPs that may be driving differences between the *S. tubifer* populations based on Weir and Cockerham F_ST_ values calculated in VCFtools v0.1.16.

### Symbiont Analyses

Five individuals were removed from the symbiont analysis due to poor quality after filtering. Trimmed and aligned bacterial sequences from the remaining 56 light organ symbiont populations were assembled with metaSPAdes v3.15.5 (Nurk et al. 2017). Prokka v1.14.6 (https://github.com/tseemann/prokka) was then used to annotate the metagenome of each light organ sample. The outgroups for this analysis include *P. lucens* strains *ajapo*4.1 (PYNQ01000001.1, Kaeding et al. 2007) and *ajapo*5.5 (GCA_030685555.1, Gould & Henderson 2023) and *P. leiognathi* strains *ljone*.10.1 (ASM3071696v1) and *lrivu*.4.1 (ASM50920v1), all of which are bioluminescent symbionts of other fish hosts. Roary v3.13.0 (Page et al. 2015) was used to run a pangenome analysis of the symbionts using the Prokka annotations, and an alignment of the core genes present in at least 95% of the samples was created. A maximum likelihood tree was then inferred with 1000 SH-aLRT and ultra-fast bootstrap replicates using the substitution model GTR+F+R4 based on the lowest BIC score in IQ-TREE v2.0.3.

To identify patterns of genetic differentiation between the bacterial symbiont populations, we aligned the trimmed reads from each light organ to the bacterial reference, *P. mandapamensis* (ASM3068539v1), using BWA-MEM v0.7.17. The aligned reads were then used as input for snippy v4.6.0 (https://github.com/tseemann/snippy), which called consensus SNPs for each light organ symbiont population using the minimum number of reads covering a site (--mincov 10) and minimum proportion of reads that differ (--minfrac 0.9) flags. Snippy-core was implemented to produce a core alignment of the consensus SNPs across all individuals, which was used as input in IQ-TREE v2.0.3 to create a phylogenetic tree using the maximum likelihood model TVM+F+R7 with the lowest BIC score including SH-aLRT and ultra-fast bootstrap based on 1000 replicates.

### Co-phylogeny Analysis

To determine whether there was any phylogenetic congruence between *Siphamia* hosts and their bioluminescent symbiont *Photobacterium mandapamensis*, the R package paco v0.4.2 (Balbuena et al. 2013; Hutchinson et al. 2017) was used to carry out a Procrustean Analysis of Cophylogeny (PACo), a global fit method based on Procrustean superimposition to assess the degree to which host-symbiont associations may contribute to co-phylogeny. The R package ape v5.7-1was also used to run a parafit analysis, which tests the degree of coevolution between hosts and symbionts. For *Siphamia*, we used the phylogeny inferred from the mitochondrial sequences described above, and we used the tree generated from SNP data for the symbiont. Both trees were altered into a dendrogram for visual representation using the R package dendextend v1.17.1 (Galili 2015).

## RESULTS

### Sequence Data

Whole genome sequencing of the light organs resulted in an average of 92,743,381 (min = 55,591,110, max = 140,448,218) raw reads per sample, while post-filtering had 87,431,907 (min = 51,440,137, max = 135,309,341) reads. On average, the host and bacteria accounted for 78.3% (SE +/-0.994%) and 21.7% (SE +/-0.994%) of the reads, respectively. The average depth coverage for the fish was 12.9, while the average depth coverage for bacteria was 983.6 (Supplementary Table 1).

### Host Analyses

Phylogenies were inferred using maximum likelihood for both whole mitochondrial genomes of the hosts sequenced in this study (n = 57), and for COI gene sequences (n = 88) comprised of both our samples and 28 additional COI sequences that were publicly available representing other *Siphamia* species (Supplementary Table 1). Both phylogenetic analyses confirmed that three distinct species were examined in this study: *S. mossambica, S. tubifer*, and *S. fuscolineata*. They also both recovered *S. mossambica* as basal to a clade containing sister taxa *S. tubifer* and *S. fuscolineata* (Figure 2). Within *S. tubifer*, there was a clade comprised only of individuals from Japan, suggesting divergence between individuals sampled from the two locations.

**Figure 1.**
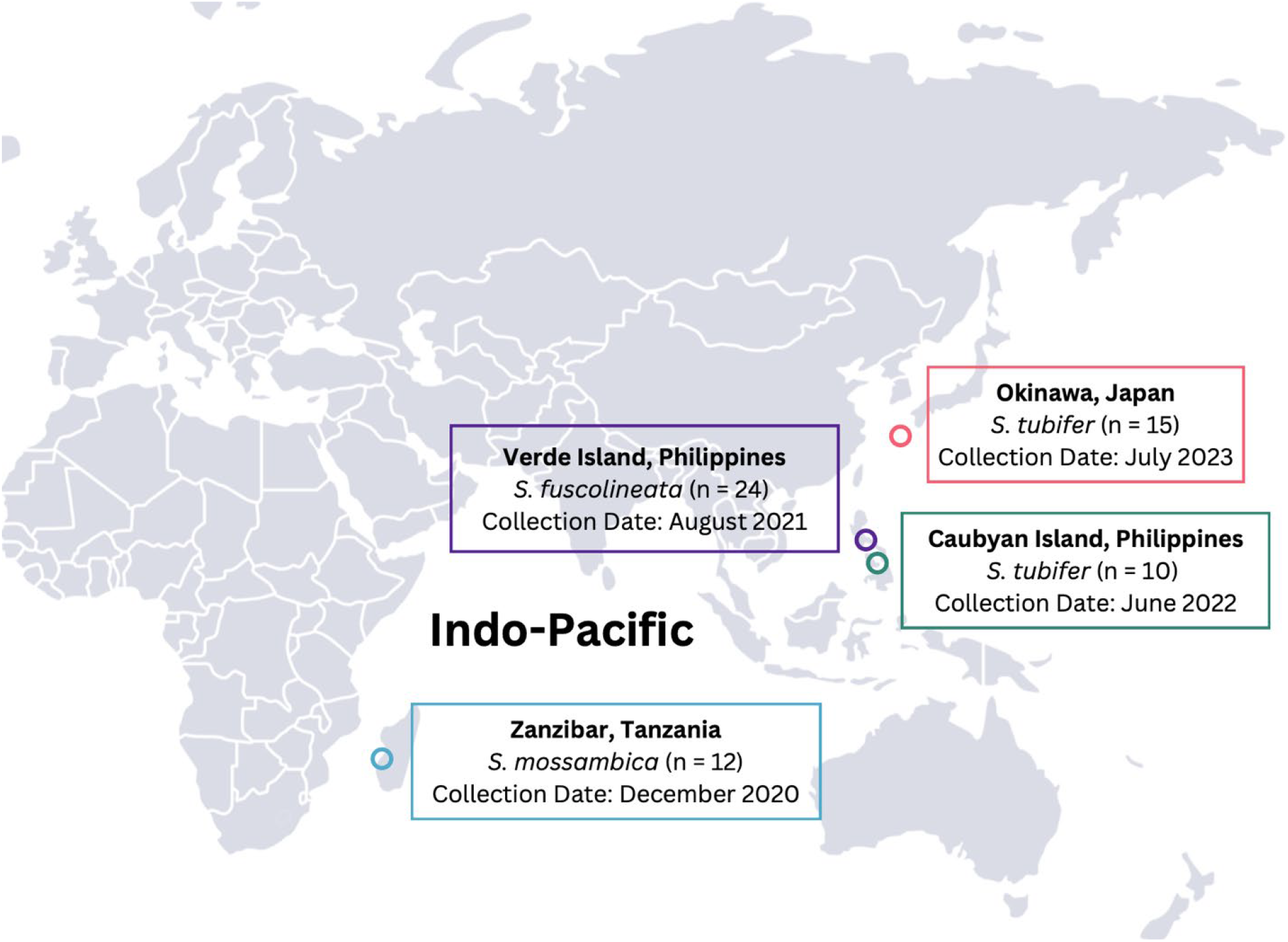
Map of collection sites for *Siphamia* species across the Indo-Pacific. Collection dates and sample sizes are indicated in the boxes.

**Figure 2.**
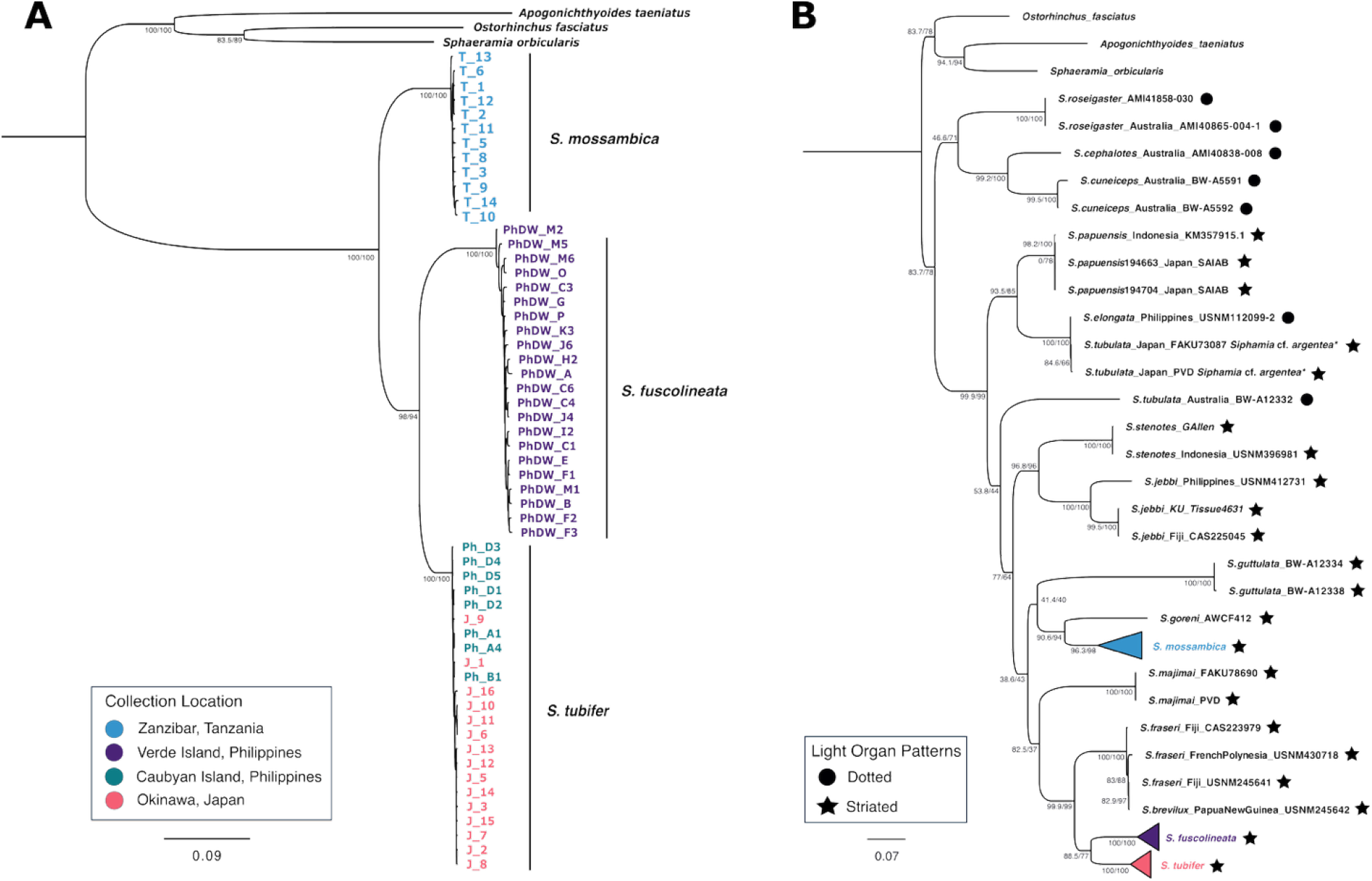
Maximum likelihood phylogenetic trees of *Siphamia species*. 2a) Phylogeny of *Siphamia* specimens sampled in this study with a substitution model of TPM2+F+R5. 2b) Phylogeny of *Siphamia species* (n=88) based on COI sequences using the TPM2+F+I+G4 substitution model. The type of light organ for each species is indicated. The three outgroups include the striped cardinalfish, *Apogonichthyoides taeniatus* (MN699562.1), the broad-banded cardinalfish, *Ostorhinchus fasciatus* (NC_058293.1) and the polka-dot cardinalfish, *Sphaeramia orbicularis* (AP018927.1). Species with an asterisk (*) are misidentified and are most likely *S. argentea*. Both figures depict SH-aLRT (%) and ultrafast bootstrap support (%) at major nodes.

The phylogeny based solely on the COI gene helped to resolve the evolutionary relationships among species in the *Siphamia* genus, particularly within the *tubifer* group. Within this tree, *S. mossambica* was sister to *S. goreni*. There was also strong support for a *tubifer* clade comprised of species with striated light organs that includes *S. tubifer*. However, due to limited sequence availability, the *tubulata* group was not clearly defined. The single *S. tubulata* specimen was placed within the *tubifer* clade with relatively low support. However, the phylogeny did show strong support for a basal clade comprised of three Australian species, all of which have spotted light organs.

A genome-wide analysis identified a total of 15,506 SNPs across the 57 individual *Siphamia* hosts. A PCA of the SNPs showed clear distinctions between the three species as confirmed by a permutational multivariate analysis of variance using distance matrices (PERMOVA) (*R*^2^ = 0.57, *F* = 23.44, Df = 3, *p* = 0.001). (Supplementary Figure 1). PC1 described 46.14% of the variation and primarily accounted for the differences between *S. mossambica* and the other two species, while PC2 explained 17.82% of the variation and accounted for the differences between *S. fuscolineata* and *S. tubifer*. Genetic variation dropped to 0.94% and 0.92% for PC3 and PC4, respectively. The two *S. tubifer* populations did not show variation when compared at this species-level scale.

An analysis of only the *S. tubifer* sequences identified 1,098,423 SNPs across all 23 individuals from Japan and the Philippines (Figure 3). A PCA based on these SNPs indicate genetic differentiation between *S. tubifer* collected from Japan and the Philippines, as confirmed by a PERMOVA (*R*^2^ = 0.077, *F* = 1.74, Df = 1, *p* = 0.001), although there was higher variance in the Philippines population. PC1 accounted for 8.11% of the variation in the data, while PC2, PC3, and PC4 accounted for 7.14%, 5.43%, and 4.69%, respectively. PC1 largely explained the difference between locations, while PC2 and PC3 captured most of the variation in the Philippines population. PC4 began to explain the variation within the population from Japan. An analysis of genetic differentiation between the two populations revealed a pairwise F_ST_ value of 0.043, however, the distribution of F_ST_ values of individual SNPs showed no clear region in the genome driving this difference (Figure 4). Additional population-level statistics were carried out to assess and compare the diversity and structure of each population (Table 1).

**Table 1.** Population statistics for S. tubifer based on SNP genotypes.

| Location | n | Ho | He | GC% | A% | $\pi$ |
| --- | --- | --- | --- | --- | --- | --- |
| Japan | 15 | 0.2973 | 0.2940 | 0.7596 | 98.36 | 0.0251 |
| Philippines | 8 | 0.2995 | 0.2933 | 0.7674 | 91.76 | 0.0475 |

Number of individual in a population (n), observed heterozygosity ( $H_o$ ), expected heterozygosity ( $H_e$ ), genotype concordance percentage (GC), percentage of total alleles observed ( $A_o$ ), nucleotide diversity ( $\pi$ ).

**Figure 3.**
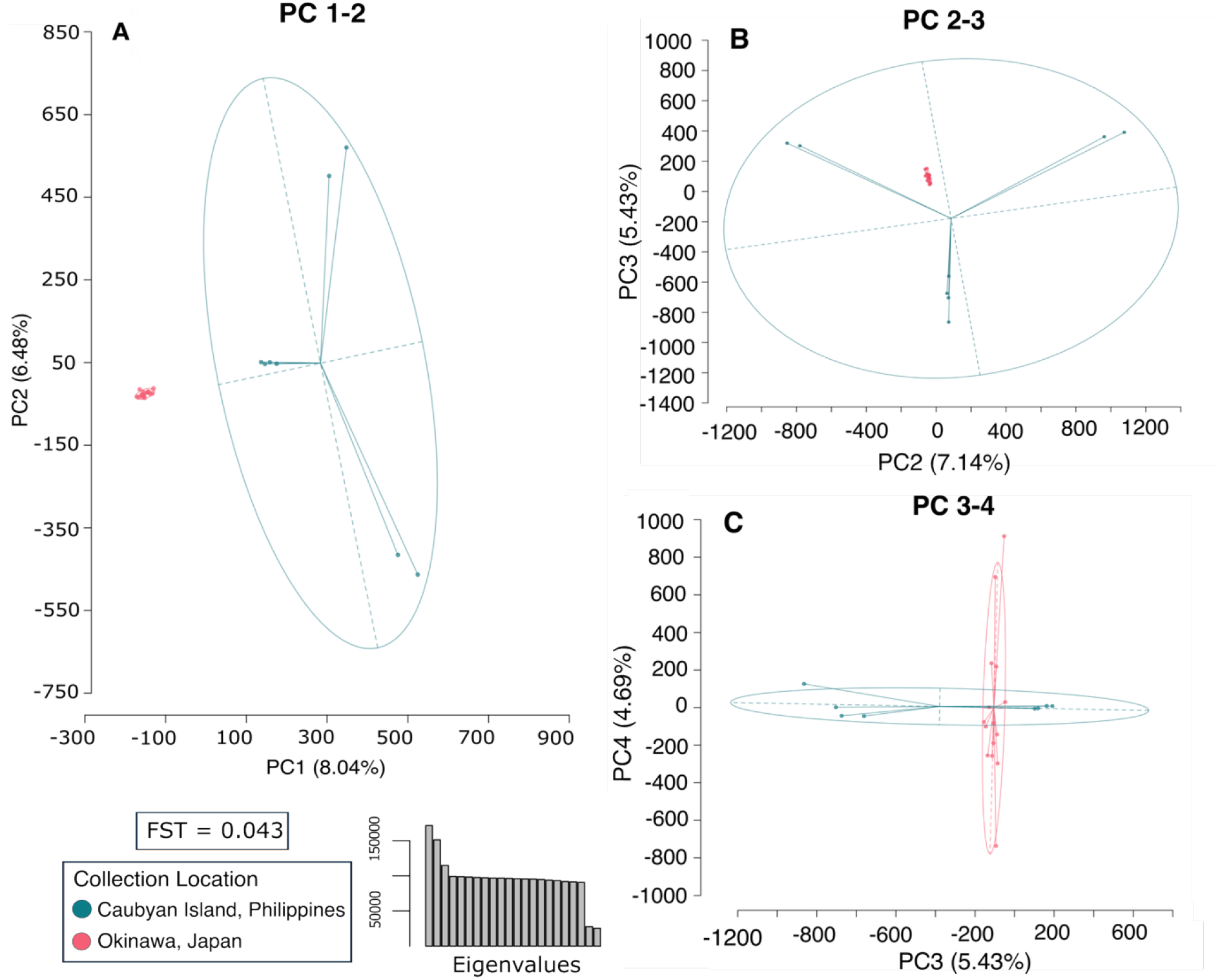
Principal Component Analysis of *Siphamia tubifer* from Japan (n = 15) and the Philippines (n = 8) shows evidence of genetic differentiation based on 1,098,423 SNPs. Plots a-c show the results for PCs 1-4. Eigenvalues representing PC1–22 are shown in the barplot, and the pairwise F_ST_ value between the two populations is included.

**Figure 4.**
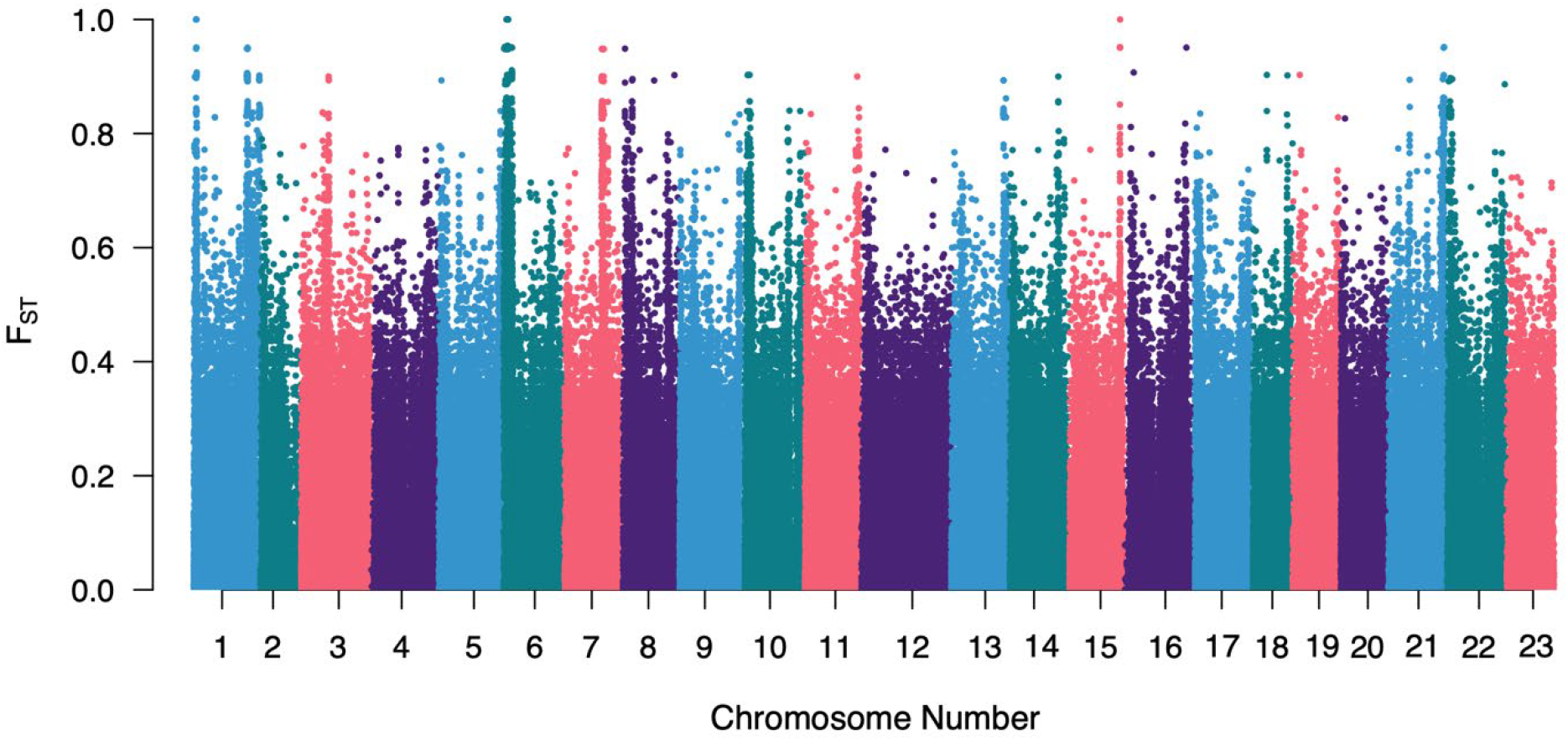
F_ST_ values of individual SNPs identified between *S. tubifer* populations from Japan and the Philippines. The location of each SNP is represented by its location in each of the corresponding chromosomes numbered along the x-axis.

**Figure 4.**
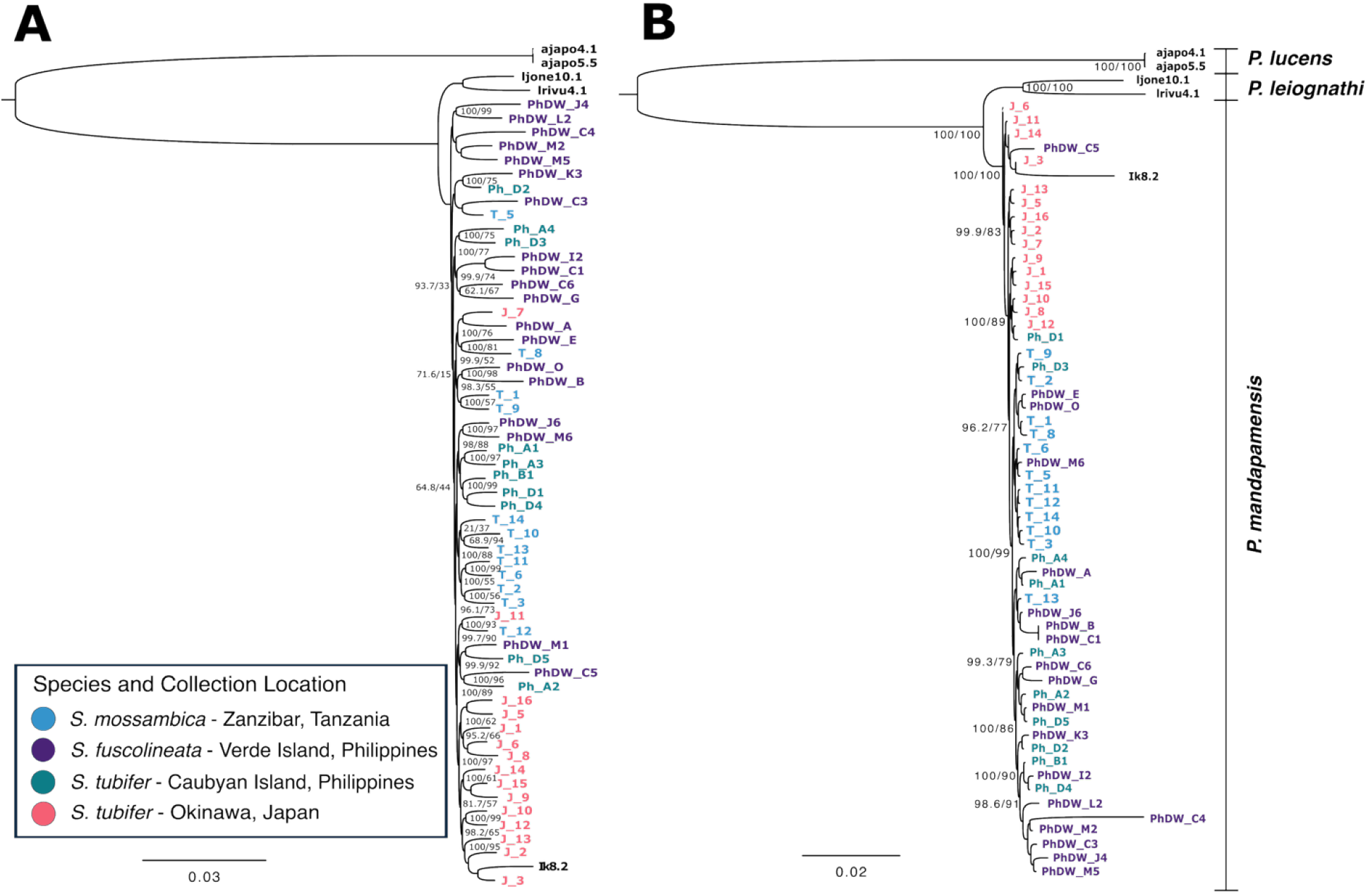
Maximum likelihood phylogeny of *Siphamia* light organ symbionts. 4a) Phylogeny is based on 2,224 core genes. Model of substitution is GTR+F+R4. Reference bacterial strains of *Photobacterium lucens* (*ajapo4*.*1, ajapo5*.*5*), *P. leiognathi* (*ljone10*.*1, lrivu4*.*1*) and *P. mandapamensis* (*Ik8*.*2)* were also included. Nodes missing SH-aLRT/bootstrap values signifies 100/100 support. 4b) Phylogeny based on 32,405 consensus SNPs. TVM+F+R7 was used as the model of substitution to determine support values. Bacterial strains of *Photobacterium lucens* (*ajapo*4.1, *ajapo*5.5), *P. leiognathi* (*ljone*10.1, *lrivu*4.1) and *P. mandapamensis* (*Ik8*.*2*) were included in the analysis as references. SH-aLRT/bootstrap values are indicated at the major nodes.

### Symbiont Analyses

#### Phylogenetics of the Symbiont

A total of 2,224 core genes were identified across the symbiont samples and used to infer a maximum likelihood phylogeny (Figure 4a). Reference strains of *Photobacterium lucens* (*ajapo4*.*1, ajapo5*.*5*), *Photobacterium leiognathi* (*ljone*10.1, *lrivu*4.1), *and P. mandapamensis* (*Ik8*.*2*) were included in the analysis to confirm the identity of the light organ symbionts based on their placement within the phylogeny. All light organ symbionts were assigned within the *P. mandapamensis* clade with strong support (100/100). Symbionts from Japan and Tanzania also formed tight clades based on their respective locations, while the Philippines symbionts were intermixed despite differences in host species, collection depth, and location.

A maximum likelihood phylogeny based on 32,405 consensus SNPs also depicted patterns of genetic variability in the symbiont associated with geography. The inferred phylogeny revealed that symbionts largely clustered according to their geographic origin, with symbionts from the same sampling location exhibiting greater genetic relatedness to one another (Figure 4b). In concordance with the core gene phylogeny, there were several distinct clades comprised exclusively of symbionts from each location, with most of the symbionts from the Philippines interspersed irrespective of which host it originated, *S. fuscolineata* or *S. tubifer*.

#### Symbiont Pangenome Analysis

To identify genetic differences between the light organ symbionts of the different hosts, we also performed a pangenome analysis based on the presence and absence of genes within each light organ metapopulation (Supplementary Figure 2). Across all light organs examined, there were 2,224 core genes (present in at least 95% of the light organs), 4,086 shell genes (present in 15% to 94% of the light organs), and 59,207 cloud genes (found in fewer than 15% of the light organs). Of these, 37,476 were singletons, present within only a single light organ, while 976 genes were present in all samples. Although the core gene phylogeny contained clades comprised of symbionts from specific locations, there were no distinguishable patterns in gene content of the symbionts that corresponded to these locations or to their host species.

### Co-phylogenetic Analysis

An analysis of the topologies of the phylogenies of the *Siphamia* hosts and their bioluminescent symbiont, *P. mandapamensis*, indicated no evidence of codivergence (Figure 5). The supporting PACo analysis showed varying co-divergence signals based on species (Supplementary Material Table 2). Co-divergence was most strongly supported by *S. tubifer* populations, in contrast to *S. mossambica*, which exhibited the weakest support. The parafit global test encapsulating all species provided no support for patterns of codivergence (p = 0.92) based on 1000 permutations.

**Figure 5.**
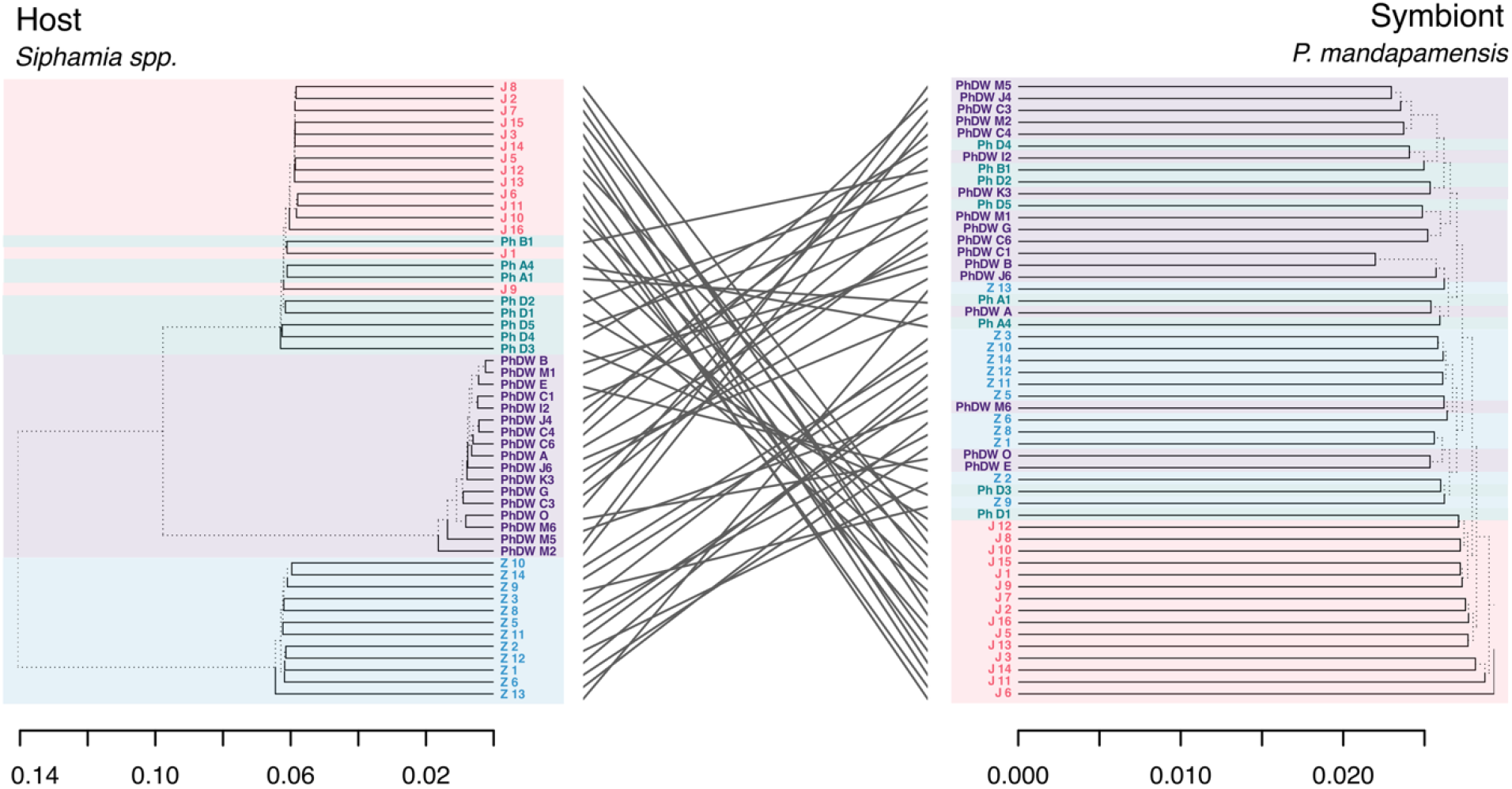
Tanglegram showing 52 host-symbiont links between *Siphamia* spp. and *Photoabcterium mandapamensis*. The host phylogeny (left) is based on whole mitochondrial genome sequences. The symbiont phylogeny (right) is based on 32,405 consensus SNPs from each corresponding light organ.

## DISCUSSION

Determining how marine organisms maintain specific partnerships with horizontally acquired microbial symbionts remains a key question in symbiosis research. While the high connectivity of ocean environments combined with the dispersal potential of marine bacteria should predict relaxed partner selection and symbiont switching, many marine organisms maintain some degree of specificity with their microbial partners (Troussellier et al. 2017). How such specificity is maintained requires examining both the evolutionary relationships between host and symbiont and the ecological mechanisms that govern symbiont acquisition. The *Siphamia*-*Photobacterium* association offers a unique opportunity to investigate these dynamics, combining high specificity across diverse hosts with broad geographic distributions and habitat dependencies that could influence both host dispersal and symbiont availability. Our whole genomic analysis of three *Siphamia* species in the *tubifer* group and their symbionts reveals how geographic, ecological and evolutionary factors interact to shape the specificity of this facultative marine symbiosis.

To understand how the bioluminescent symbiosis has evolved and persisted, we first examined the evolutionary relationships among hosts and their biogeographic distributions. Our phylogenetic analysis indicates that *S. mossambica* is basal to both *S. fuscolineata* and *S. tubifer*, which are closely related sister species. Incorporating additional *Siphamia* species in the COI gene analysis supported that these species all belong to the more recently derived *tubifer* group. Within the *tubifer* group, *S. tubifer* has the largest distribution, overlapping in range with both *S. mossambica* and *S. fuscolineata* (Gon & Allen 2012). In comparison, *S. mossambica* is endemic to the Western Indian Ocean, has a much narrower distribution, and is geographically isolated from the other species examined. Members of the *tubifer* group typically associate with sea urchin (or other echinoderm) hosts throughout the Indo-Pacific (Gon & Allen 2012). All three *Siphamia* species in this study formed associations with urchins, but each was collected from a different species. *Siphamia tubifer* preferred the long spined sea urchins (*Diadema setosum*), whereas *S. fuscolineata* were collected with fire urchins (*Astropyga radiata*) at greater depths (>30m) than the others. This difference in urchin use and depth could be a result of niche partitioning leading to the divergence of *S. tubifer* and *S. fuscolineata*.

In contrast, members of the *tubulata* group, the less speciose of the two groups, exhibit broader substrate use, including corals, sea grass, sand and rubble (Gon & Allen 2012). The evolutionary history of habitat diversification across the genus remains poorly understood, in part due to the paucity of genetic data available for species in the *tubulata* group. Although relatively little is known about these species, they share a dotted light organ pattern with the basal clade of Australian species, suggesting this is likely the ancestral state, while the striated pattern in the *tubifer* group is a derived trait. The functional significance of these different color patterns to the symbiosis remains unknown. Additional research on the *tubulata* group and their bioluminescent symbionts is needed to develop an understanding of the evolutionary history and potential drivers of speciation within the group as well as the specificity of the symbiosis.

In addition to characterizing genetic divergence among *Siphamia* species, we analyzed population-level differences within the most widespread *Siphamia* species. The two populations of *S. tubifer* examined were collected from regions that are highly influenced by the Kuroshio Current, a major ocean current that flows from the eastern edge of the Philippines to the eastern coasts of Japan (Barkley 1973). Despite the influence of the Kuroshio Current, populations of *S. tubifer* from Japan and the Philippines are genetically differentiated from one another, as seen for other reef fish in the region (Ackiss et al. 2018, Islam et al. 2022, Roberts 2022). However, a previous study examining patterns of spatial and temporal genetic variation in *S. tubifer* within the Okinawan Islands found no evidence of genetic differentiation at this finer scale (Gould & Dunlap 2017). Our results also suggest that the Philippines population had a higher level of genetic variability compared to the Japanese population. Future studies incorporating broader geographic sampling and larger population sizes of *S. tubifer* along the Kuroshio Current would help reveal the evolutionary mechanisms driving population connectivity patterns in this widespread species.

In parallel to our host analysis, our genomic analysis of the symbiont confirms that the specificity of the association is maintained across multiple host species and throughout the Indo-Pacific. All three host species formed strict symbiotic partnerships with *P. mandapamensis*, a subspecies of *P. leiognathi*, supporting previous results suggesting that the symbiosis is highly conserved throughout the *Siphamia* genus (Gould et al. 2021). In comparison, most symbiotically luminous fish and squid species do not exhibit this degree of specificity (Dunlap et al. 2007). There have even been instances of co-symbiosis of *P. leiognathi* and *P. mandapamensis* within the same light organ of several fish species (Kaeding et al. 2007). It is still largely unknown why or how *Siphamia* recognize and strictly select *P. mandapamensis* from the environment, and whether the fish can distinguish *P. mandapamensis* from closely related *P. leiognathi*.

The exceptional degree of specificity observed across *Siphamia* hosts poses interesting questions about the mechanisms involved in such precise partner recognition. *Photobacterium leiognathi* and *P. mandapamensis* are found in overlapping regions across the Indo-Pacific, and both species are facultative symbionts of luminous fishes. They have high overall genetic similarity (96.5% shared average nucleotide identities, Gould & Henderson 2023) and are indistinguishable at the 16S rRNA gene, leading to changes in taxonomic classification over the years (Boisvert et al. 1967; Hendrie et al. 1970; Reichelt and Baumann 1975; Urbanczyk et al. 2013). A comparison of the luminescence genes between *P. mandapamensis* and *P. leiognathi* revealed differences in the *lux* operon, distinguished by the presence of the *luxF* gene in *P. mandapamensis* and its absence in *P. leiognathi* (Ast and Dunlap 2004). Furthermore, both strains differ in their type II secretion systems (Gould and Henderson 2023). It is unclear if these differences are useful for hosts to distinguish between the two symbiont species. Determining whether these factors influence the hosts’ ability to identify and select *P. mandapamensis* from the environment will provide insight into the molecular mechanisms involved in regulating the specificity of the association.

In addition to the genetic mechanisms involved in regulating specificity, the host’s behavior and ecology may play an important role. *Siphamia* hosts shed excess symbionts back into the free-living environment with fecal waste (Dunlap & Nakamura 2011), which could increase the local density of *P. mandapamensis* in the surrounding seawater. Settling *Siphamia* larvae would have a higher chance of encountering *P. mandapamensis* near populations of adult fish as opposed to when they are in the open ocean, where the relative abundance of *Photobacterium* spp. is low (Troussellier et al. 2017). This local enrichment would also help explain the biogeographic patterns of symbiotic *P. mandapamensis* observed in this and other studies (Gould and Dunlap 2019; Gould et al. 2023). Future research on the influence of the host fish on free-living symbiont populations will improve our understanding of the role local enrichment plays in symbiont acquisition and the specificity of the association.

While these ecological mechanisms may promote specificity, our genomic analysis of the symbiont tells a more complex story. Symbionts from Tanzania, the Philippines, and Japan largely form their own clades, yet a comparative analysis of the symbiont pangenomes revealed no discernable patterns in gene content between hosts from these locations. This lack of functional differentiation may reflect the limitations of our whole light organ sequencing approach, which analyzed bacterial symbionts based on the presence and absence of genes within an entire light organ rather than examining differences between individual symbiont genomes. Analyzing genomes at the isolate level might provide the necessary resolution to uncover specific genetic differences between symbionts from different locations or host species. Interestingly, the light organ symbionts from two hosts (*S. tubifer* and *S. fuscolineata*) collected at varying depths (approximately 3 and 30 meters) in the Philippines showed no evidence of phylogenetic divergence. This could indicate that larvae from both hosts either acquired their bioluminescent symbiont from similar environmental pools of bacteria, or that local currents promote dispersal and mixing of marine bacteria in the region.

Despite the geographic structuring observed in the symbiont, a comparative phylogenetic analysis revealed no evidence of co-divergence between *Siphamia* hosts and their bioluminescent symbionts. This is congruent with that of other symbiotically luminous fishes, for which co-divergence has not been observed (Dunlap et al. 2007). The previous study used COI and 16S rRNA focusing on a much broader phylogenetic range of hosts and symbionts, whereas our research focused strictly on *Siphamia* hosts in the *tubifer* group and a subspecies of a bacterium. The absence of co-divergence likely reflects the facultative nature of the symbiont. *Photobacterium mandapamensis* can persist in the free-living environment, preventing any dependency on the host for survival or reproduction (Urbanczyk et al. 2010). While co-evolution has been described in other similar, facultative associations, such as the squid-*Aliivibrio* model (Nishiguchi et al. 1998), our study revealed patterns of genetic divergence that are incongruent with host divergence. Instead, genetic structure in the symbiont appears to be driven by geographic factors, whereas host divergence likely results from other factors such as habitat partitioning discussed above.

In summary, our genomic analysis of *Siphamia* species in the *tubifer* group and their bioluminescent symbionts reveals a highly conserved association throughout the Indo-Pacific. This is the first study to characterize the light organ symbionts of *S. mossambica* and *S. fuscolineata*, confirming that along with *S. tubifer* all hosts partner exclusively with *P. mandapamensis*, a subspecies of *P. leiognathi*. As for the host, we were able to help resolve the evolutionary relationship of the *tubifer* clade of *Siphamia* and found evidence of genetic differentiation between populations of *S. tubifer* from Japan and the Philippines. We also found evidence of phylogeographic structure in the symbiont yet found no evidence of host-symbiont co-divergence. This suggests ecological processes, including local enrichment of symbionts by the host fish, rather than co-evolutionary dynamics, help promote specificity of the association. These findings challenge the paradigm in marine microbial ecology that “everything is everywhere” by demonstrating how host ecology can promote specificity in facultative symbioses despite the high dispersal potential of marine bacteria, offering new insights into how horizontally acquired microbial partnerships persist in connected marine environments.

