## Supplementary Table 1 for "Genomic Insights into a Highly Specific Marine Symbiosis Uncovers Geographic Structuring Without Co-Divergence Between *Siphamia* Cardinalfish and Their Bioluminescent Symbiont"

Supplementary Table 1*. Siphamia* individuals collected for this study. Listed is each specimen’s ID, species, collection location, collection date, approximate collection depth, and standard length (SL). One asterisk (*) represents samples used for bacteria only and two asterisks (**) indicates samples used for fish only due to poor quality reads.

| **Specimen ID** | **SL (cm)** | **Raw Reads** | **Reads after Trimming** | **Primary Fish Alignments** | **Primary Bacterial Alignments** | **SNP Fish** | **SNP Bacteria** | **Average Depth Coverage for Fish** | **Average Depth Coverage for Bacteria** |
| --- | --- | --- | --- | --- | --- | --- | --- | --- | --- |
| J_1 | 3.04 | 94137198 | 85798876 | 105167029 | 59159326 | 7113050 | 15048 | 10.9 | 1539 |
| J_2 | 3.5 | 102508800 | 92226404 | 139751700 | 25841042 | 7990097 | 18393 | 14.9 | 636 |
| J_3** | 2.63 | 83561266 | 75756198 | 102757698 | N/A | 7038308 | N/A | 10.6 | N/A |
| J_5 | 2.45 | 108079125 | 96830100 | 126760320 | 55234972 | 7559619 | 16922 | 13.5 | 1441 |
| J_6 | 2.63 | 91518696 | 83368048 | 118720509 | 35915107 | 7465207 | 13975 | 12.4 | 900 |
| J_7 | 2.58 | 112046171 | 103016912 | 140575636 | 53832970 | 7720419 | 19098 | 14.7 | 1361 |
| J_8 | 2.67 | 98924830 | 92655579 | 115358543 | 53710006 | 7330448 | 18151 | 11.9 | 1399 |
| J_9 | 2.73 | 83634912 | 80629397 | 111939057 | 41487737 | 7534926 | 16657 | 12.9 | 1155 |
| J_10 | 2.54 | 85534886 | 81790393 | 118369041 | 34343577 | 7700288 | 13193 | 13.7 | 953 |
| J_11 | 2.35 | 110527521 | 106127303 | 152408342 | 46961609 | 8088290 | 12329 | 17.4 | 1283 |
| J_12 | 2.07 | 99487242 | 95912608 | 131997022 | 50847335 | 7894524 | 13320 | 15.2 | 1421 |
| J_13 | 2.19 | 89995333 | 86339570 | 121396113 | 42427205 | 7679455 | 14950 | 13.6 | 1143 |
| J_14 | 1.82 | 97425087 | 93506813 | 133769819 | 42343822 | 7851317 | 10049 | 15.1 | 1134 |
| J_15 | 2.01 | 94495797 | 90324066 | 128610885 | 38494972 | 7797582 | 13709 | 14.8 | 1037 |
| J_16 | 1.52 | 87850775 | 83577727 | 120412069 | 33650630 | 7639240 | 15682 | 13.2 | 870 |
| Ph_A1 | 2.32 | 76161155 | 70766195 | 106257758 | 19075779 | 7428821 | 15684 | 11.3 | 465 |
| Ph_A2* | 2.51 | 122047081 | 105847176 | N/A | 132759894 | N/A | 19271 | N/A | 2850 |
| Ph_A3* | 2.25 | 96819629 | 89902000 | N/A | 106402194 | N/A | 17090 | N/A | 2561 |
| Ph_A4 | 2.34 | 102754612 | 97117232 | 73847190* | 170206100 | 6933373 | 22634 | 8.45 | 4250 |
| Ph_B | 2.4 | 89135827 | 84172377 | 117687335 | 40317918 | 7356240 | 13219 | 11.9 | 991 |
| Ph_D1 | 3.48 | 108174306 | 100583646 | 148574222 | 34847603 | 7965142 | 18377 | 14.6 | 829 |
| Ph_D2 | 2.16 | 102462501 | 97101272 | 150839507 | 22510374 | 8020939 | 14831 | 16 | 576 |
| Ph_D3 | 2.11 | 91036184 | 84877568 | 127943221 | 25728951 | 7958224 | 20157 | 13.6 | 656 |
| Ph_D4 | 2.22 | 87321374 | 81023102 | 116326519 | 26231721 | 7246495 | 17982 | 11.4 | 610 |
| Ph_D5 | 2.55 | 65482419 | 59116909 | 70474678* | 31346853 | 6785336 | 19082 | 8.06 | 746 |
| PhDW_A | 3.29 | 84449579 | 81710736 | 121764893 | 15823915 | 22197119 | 29208 | 14.2 | 398 |
| PhDW_B | 2.53 | 113803363 | 106525810 | 148685105 | 31005612 | 12717457 | 32602 | 15.8 | 759 |
| PhDW_C1 | 2.32 | 102760479 | 98841452 | 150306914 | 13373762 | 22982688 | 32602 | 17.5 | 329 |
| PhDW_C3 | 2.59 | 107745257 | 92906002 | 109175394 | 57218561 | 17785663 | 26311 | 9.3 | 1238 |
| PhDW_C4 | 2.07 | 91251600 | 84275730 | 121494976 | 18179968 | 21526086 | 69310 | 13.8 | 423 |
| PhDW_C5* | 2.51 | 68013541 | 61876592 | N/A | 67830625 | N/A | 34044 | N/A | 1734 |

| **Specimen**  **ID** | **SL (cm)** | **Raw Reads** | **Reads after Trimming** | **Primary Fish Alignments** | **Primary Bacterial Alignments** | **SNP Fish** | **SNP Bacteria** | **Average Depth Coverage for Fish** | **Average Depth Coverage for Bacteria** |
| --- | --- | --- | --- | --- | --- | --- | --- | --- | --- |
| PhDW_C6 | 2.28 | 82363207 | 79650339 | 118767911 | 15362991 | 22247805 | 23510 | 14.2 | 410 |
| PhDW_E | 2.38 | 137299558 | 131709162 | 182763590 | 44625735 | 23467440 | 19843 | 21.5 | 1192 |
| PhDW_F1** | 2.25 | 137187225 | 131365070 | 209622733 | N/A | 24267424 | N/A | 25.7 | N/A |
| PhDW_F2** | 2.31 | 87825089 | 84238662 | 134679121 | N/A | 22499271 | N/A | 16.7 | N/A |
| PhDW_F3** | 2.6 | 67034268 | 64971009 | 104095181 | N/A | 21673630 | N/A | 12.7 | N/A |
| PhDW_G | 2.21 | 55591110 | 51440137 | 76890722 | 4925180 | 18582896 | 31385 | 8.29 | 107 |
| PhDW_P** | 2.41 | 89021079 | 81844944 | 130045846 | N/A | 21789120 | N/A | 14 | N/A |
| PhDW_H2** | 2.46 | 125633616 | 117529445 | 97274012 | N/A | 19726719 | N/A | 10.8 | N/A |
| PhDW_I2 | 2.67 | 84551436 | 81068974 | 102390880 | 40931962 | 19909782 | 24345 | 10.6 | 1003 |
| PhDW_J4 | 2.05 | 105000568 | 98658685 | 98158418 | 44969932 | 18121572 | 33138 | 9 | 1060 |
| PhDW_J6 | 2.68 | 85225501 | 80576033 | 106594474 | 33178538 | 20657676 | 16274 | 11.4 | 808 |
| PhDW_K3 | 2.57 | 92556733 | 85424744 | 119369314 | 20007589 | 20621684 | 24259 | 12.2 | 439 |
| PhDW_L2* | 2.26 | 96256298 | 92442644 | N/A | 14449485 | N/A | 25453 | N/A | 285 |
| PhDW_M1 | 2.29 | 84955042 | 82260794 | 124852804 | 13540867 | 22236570 | 20078 | 14.7 | 338 |
| PhDW_M2 | 2.68 | 130608094 | 125052932 | 163233292 | 48458869 | 22764181 | 18457 | 17.9 | 1225 |
| PhDW_M5 | 2.36 | 97979272 | 91338201 | 114655808 | 37039265 | 20697646 | 18221 | 12.3 | 923 |
| PhDW_M6 | 2.16 | 140448218 | 135309341 | 199374519 | 29811657 | 23534940 | 19113 | 23.3 | 736 |
| PhDW_O | 2.55 | 86830151 | 83205575 | 114477775 | 28958165 | 20973553 | 20785 | 12 | 682 |
| T_1 | 2.55 | 68262044 | 65162716 | 83125741 | 17431711 | 28770809 | 16034 | 9.19 | 437 |
| T_2 | 2.92 | 92992800 | 89980969 | 106214466 | 41385181 | 30485201 | 16479 | 11.8 | 1098 |
| T_3 | 2.65 | 72828619 | 70848930 | 82919069 | 32105020 | 29338788 | 18344 | 9.48 | 892 |
| T_5 | 2.07 | 76556805 | 73997750 | 92045830 | 24525585 | 29907969 | 18064 | 10.5 | 646 |
| T_6 | 1.69 | 67846591 | 65949814 | 82406088 | 19681439 | 29311143 | 16528 | 9.62 | 530 |
| T_8 | 1.77 | 79377961 | 76754409 | 90747273 | 33745081 | 29663019 | 23616 | 10.3 | 926 |
| T_9 | 1.37 | 71708363 | 69650213 | 79459413 | 35470241 | 28919223 | 16745 | 9.15 | 1006 |
| T_10 | 1.54 | 72847615 | 70237706 | 81352712 | 34412612 | 6699861 | 18059 | 9 | 931 |
| T_11 | 1.61 | 68195919 | 65152624 | 80766364 | 21357015 | 27873244 | 14664 | 8.49 | 523 |
| T_12 | 1.47 | 92666145 | 89838243 | 107951796 | 38156346 | 30548950 | 15238 | 11.9 | 1017 |
| T_13 | 1.78 | 83529968 | 80238205 | 98374719 | 26125608 | 29841838 | 15643 | 10.6 | 648 |
| T_14 | 1.25 | 75020442 | 72944298 | 93164162 | 19916811 | 30548375 | 14591 | 11.3 | 551 |
