## Supplementary Table 2 for "Genomic Insights into a Highly Specific Marine Symbiosis Uncovers Geographic Structuring Without Co-Divergence Between *Siphamia* Cardinalfish and Their Bioluminescent Symbiont"

Supplementary Table 2. A procrustean approach to assess phylogenetic congruence between *Siphamia* species and the symbiont *Photobacterium mandapamensis*.

| **res** | **Relationships** |
| --- | --- |
| 0.076256 | SJ_1-J_1 |
| 0.080913 | SJ_14-J_14 |
| 0.079253 | SJ_16-J_16 |
| 0.080232 | SJ_5-J_5 |
| 0.079982 | SJ_6-J_6 |
| 0.081626 | SJ_8-J_8 |
| 0.086907 | SJ_11-J_11 |
| 0.152174 | SPhDW_B-PhDW_B |
| 0.088383 | SJ_7-J_7 |
| 0.151325 | SPhDW_A-PhDW_A |
| 0.135129 | SPhDW_I2-PhDW_I2 |
| 0.135747 | SPhDW_C1-PhDW_C1 |
| 0.136914 | SPhDW_G-PhDW_G |
| 0.139937 | SPhDW_K3-PhDW_K3 |
| 0.090343 | SPh_D2-Ph_D2 |
| 0.139524 | SPhDW_C3-PhDW_C3 |
| 0.196614 | SZ_5-Z_5 |
| 0.143196 | SPhDW_J4-PhDW_J4 |
| 0.123946 | SPhDW_M2-PhDW_M2 |
| 0.128807 | SPhDW_M5-PhDW_M5 |
| 0.141088 | SPhDW_C4-PhDW_C4 |
| 0.154858 | SPhDW_M1-PhDW_M1 |
| 0.187934 | SZ_1-Z_1 |
| 0.186849 | SZ_8-Z_8 |
| 0.191207 | SZ_9-Z_9 |
| 0.147617 | SPhDW_C6-PhDW_C6 |
| 0.149557 | SPhDW_E-PhDW_E |
| 0.192738 | SZ_10-Z_10 |
| 0.185013 | SZ_13-Z_13 |
| 0.186111 | SZ_11-Z_11 |
| 0.185709 | SZ_6-Z_6 |
| 0.185784 | SZ_2-Z_2 |
| 0.186672 | SZ_3-Z_3 |
| 0.193331 | SZ_14-Z_14 |
| 0.149282 | SPhDW_J6-PhDW_J6 |
| 0.19687 | SZ_12-Z_12 |
| 0.141003 | SPhDW_M6-PhDW_M6 |
| 0.083036 | SPh_A1-Ph_A1 |
| 0.140745 | SPhDW_O-PhDW_O |
| 0.082131 | SPh_B1-Ph_B1 |
| 0.083052 | SPh_D1-Ph_D1 |
| 0.079403 | SPh_D4-Ph_D4 |
| 0.079891 | SJ_10-J_10 |
| 0.079412 | SJ_12-J_12 |
| 0.079164 | SJ_15-J_15 |
| 0.074239 | SJ_9-J_9 |
| 0.079321 | SJ_13-J_13 |
| 0.07952 | SJ_2-J_2 |
| 0.078445 | SJ_3-J_3 |

Residuals (res) are raw Procrustean residuals of each link and displays interaction-specific cophylogenetic contributions. Pairs with lower residuals show stronger co-divergence signals, whereas pairs with higher residuals show weaker co-divergence signals. The host names are indicated with an ‘S’ at the beginning of the sample name.
