## Supplementary figures and images for "Genomic Insights into a Highly Specific Marine Symbiosis Uncovers Geographic Structuring Without Co-Divergence Between *Siphamia* Cardinalfish and Their Bioluminescent Symbiont"

### Supplementary Figure 2

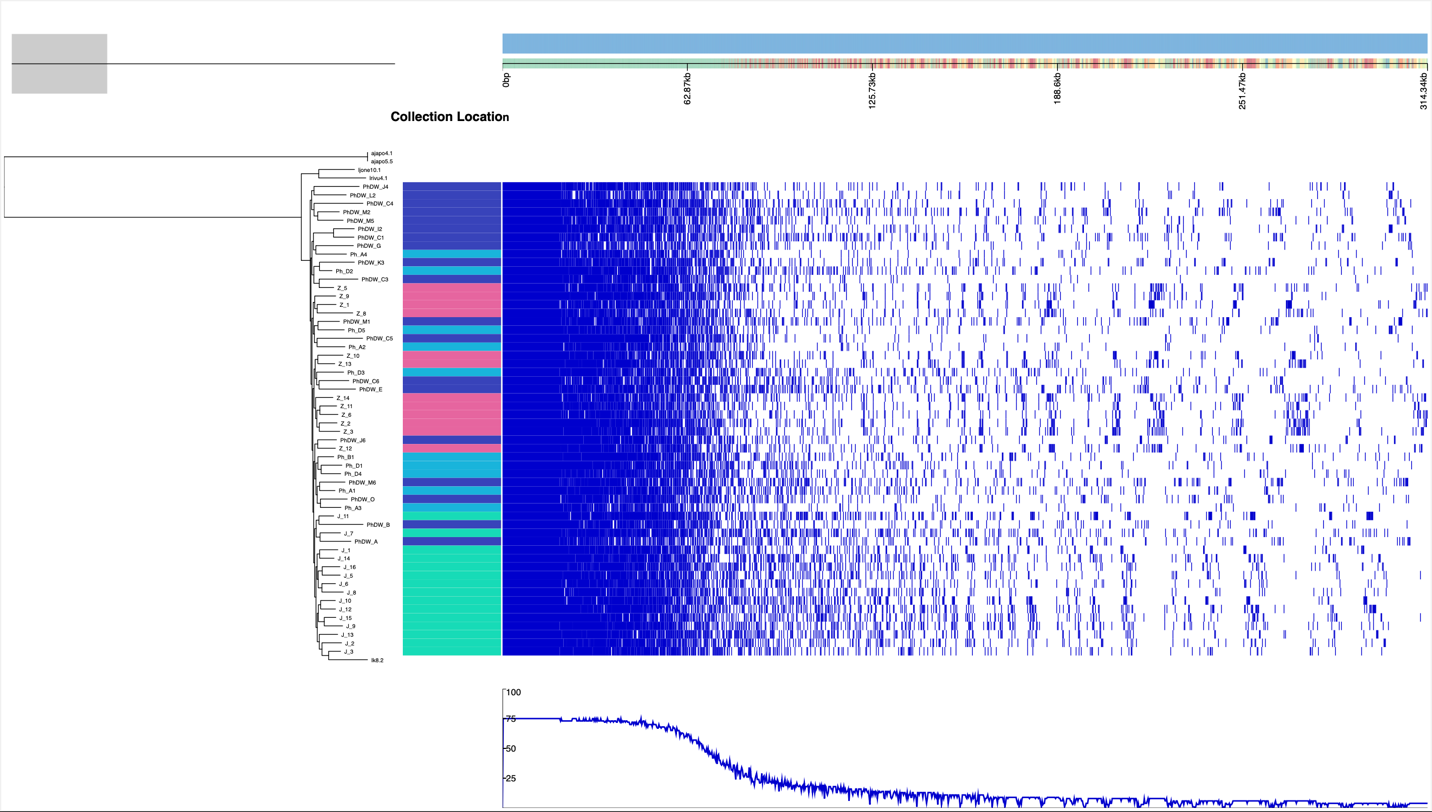
Supplementary Figure 2. A pangenome analysis of *Photobacterium mandapamensis*.
